# RiboZAP: Custom probe design for rRNA depletion in complex metatranscriptomes

**DOI:** 10.1101/2025.11.07.687087

**Authors:** Samuel Bunga, Asako Tan, Morgan Roos, Scott Kuersten

## Abstract

Metatranscriptomic (MetaT) sequencing provides critical insights into the gene expression and functional activity of microbial communities. However, its utility is limited by the overwhelming abundance of ribosomal RNA (rRNA), which typically represents ≥90% of total RNA [1-2]. A major obstacle to efficient MetaT analysis is the removal of highly abundant rRNA transcripts present in complex microbial communities, which may contain thousands of species. Although commercial rRNA depletion kits can effectively reduce rRNA content, they are typically optimized for specific host microbiomes and often underperform in others. For example, probes designed for the human gut microbiome frequently show reduced efficiency when applied to non-human samples such as mouse cecal donor samples - a common model in microbiome research. Regardless of the depletion strategy used, designing rRNA removal probes solely based on a microbiome’s taxonomic composition often requires an extensive number of probes, making the approach expensive, difficult to manufacture, and sometimes technically impractical [3]. Here, we present RiboZAP, a species-agnostic computational pipeline for designing custom RNase H depletion probes directly from MetaT sequencing data and without prior knowledge of sample composition. Our results show that the probes generated with RiboZAP are efficient for removal of rRNA content, increasing messenger RNA (mRNA), and improving transcriptome coverage. This provides a cost-effective approach to maximize the value of MetaT sequencing.

## Background

Over the past decade, rapid advances in next-generation sequencing (NGS) have revolutionized microbiome research. Higher throughput, lower costs, and improved accuracy have enabled researchers to explore microbial communities from a wide range of environments with unprecedented depth and resolution [4]. Both 16S rRNA gene sequencing and shotgun metagenomics (MetaG) are widely used to investigate microbial communities across diverse environments. The 16S approach provides a cost-effective means of profiling community composition, but it offers limited taxonomic resolution and no direct insights into functional potential [5]. By contrast, MetaG profiles all DNA present in a sample, supporting higher-resolution taxonomic profiling, often at the genus or species level [6]. While these approaches have been invaluable for mapping microbial diversity and identifying rare or novel taxa, they capture only the genetic profile of microbial communities. To understand the dynamic nature of the microbiome, it is useful to determine which genes are actively expressed and which organisms are metabolically active at the time of sampling. Such insights are critical for linking microbial presence to ecological function. MetaT addresses this gap by sequencing the community’s RNA content, enabling measurement of gene expression, metabolic pathway activity, and host-microbe interactions under different conditions. A major challenge in MetaT sequencing is the overwhelming abundance of rRNA transcripts, which can consume ≥90% of reads yet add no value to functional profiling because rRNA is a conserved structural RNA rather than mRNA, so it dilutes coverage of low-abundance transcripts, inflates sequencing cost, and reduces sensitivity to pathway-level signals. As a result, sequencing total RNA without rRNA depletion leaves only a small percentage of reads mapping to mRNA, wasting depth and reducing sensitivity. Efficient rRNA depletion is therefore essential for MetaT.

A commonly used strategy for rRNA depletion is enzymatic probe-based targeting. In this approach, antisense oligonucleotides are designed to hybridize specifically to rRNA sequences within the RNA pool. The resulting RNA-DNA hybrids are then treated with RNase H, effectively removing the targeted rRNA [7]. Commercially available kits, such as Ribo-Zero Plus (Illumina) perform effectively in human samples. However, because these kits are primarily designed for specific hosts, they are less efficient in complex microbiome samples where microbial rRNA originates from many different organisms with highly divergent sequences.

In a previous study, we introduced a rational probe design strategy in which human gut microbiome sequencing data were analyzed to identify and prioritize the most abundant rRNA sequences [7]. The resulting probe set enabled efficient enzymatic depletion of rRNA from both adult and infant stool samples with minimal bias. This approach achieved robust rRNA removal across multiple human microbiome sample types, including stool, tongue, and vaginal swabs, with more than 60% of sequencing reads retained for MetaT analysis. This method has since been commercialized as the Ribo-Zero Plus Microbiome (RZPM) kit. However, when we evaluated RZPM on mouse cecal MetaT samples, depletion efficiency was reduced, likely due to insufficient probe content specifically tuned to deplete rRNA in mouse microbiome sample types. To address this gap, we designed additional probes and combined them with RZPM, which proved to be effective in retaining ∼75% mRNA-richreads for MetaT analysis. We reported these results previously [3].

To make this approach broadly accessible and reproducible for other sample types, we developed RiboZAP, a standalone, open-source application. RiboZAP automates the entire custom probe design process, from identifying abundant rRNA sequences in MetaT data to generating optimized probe sets, making it adaptable to any microbiome of interest. The pipeline also provides users with an estimate of expected depletion efficiency prior to laboratory synthesis, enabling cost-effective optimization of probe sets for diverse microbial communities.

### Implementation

Motivated by the growing preference for modular, containerized workflows that maximize ease of installation, reproducibility, maintainability, and portability across broad range of computing environments, we implemented RiboZAP using the Nextflow workflow language v25.04.2 [18] and distributed it as a Docker image to have a consistent runtime. The container is built from Miniconda3 and bundles Python 3.12, Biopython 1.85 [13], BEDTools 2.31.1 [14], BLAST+ 2.16.0 [17], SAMtools 1.21 [19], MultiQC 1.31 [15], and SortMeRNA 4.3.6 [20]. Curated SortMeRNA rRNA databases are downloaded and indexed during the image build. The packaging and orchestration for RiboZAP is distributed as a Python package exposing a single command line interface (CLI) entry point. The CLI acts as a thin orchestration layer: it validates inputs/parameters, then invokes the RiboZAP Docker image and launches the embedded Nextflow pipeline inside the container, passing configuration at runtime. This design removes local dependency management (only Docker is required), fixed software versions, and reproduceable behavior on any UNIX-like computing environment. All key parameters (e.g., coverage thresholds, probe length, tiling gap) are user configurable. Detailed usage and examples are provided in supplementary file 1 and the project README.

The pipeline consists of two subworkflows:

1. probe design, which identifies high abundance rRNA regions from MetaT reads and generates non-redundant RNase H probe sets.
2. in silico validation, which evaluates designed probes against reference rRNA to estimate depletion efficiency prior to synthesis.

### Input and configurations

RiboZAP accepts FASTQ reads, either single-end or paired-end. However, if the intention is to run RiboZAP specifically on rRNA reads, we recommend using a k-mer–based trimmer such as BBDuk [16] to extract rRNA-matching reads into separate FASTQ files before running the tool. Inputs are supplied via a CSV sample sheet with columns sample_id and read1, and read2 if paired-end; absolute paths are recommended. Curated rRNA references (SILVA and Rfam) and a prebuilt aggregate FASTA for coverage calculations are bundled in the container and resolved at runtime, so no external downloads are required.

RiboZAP is designed to run on UNIX-like systems and scales from a personal computer to an HPC cluster as users can set the resources like CPU and memory limits. For instance, setting “--cpus 4 --memory 16” on a laptop, or higher values on a large machine to shorten runtime. Any interrupted analyses can be continued with passing a --resume flag as it reuses completed work and continues from the last successful step instead of starting over. Configurable probe design parameters include the coverage threshold “--coverage-threshold” that defines merged high-coverage blocks (default ≥500), the number of top regions “--num-cov-regions” retained per sample (default 50), probe tiling gap “--probe-tiling-gap” (default 25-nt), “--probe-length” (default 50), can be set to any value between 25-nt and 100-nt, and “--padding” option extends each alignment by 50 bp on both sides for *in silico* testing.

**Figure 1.**
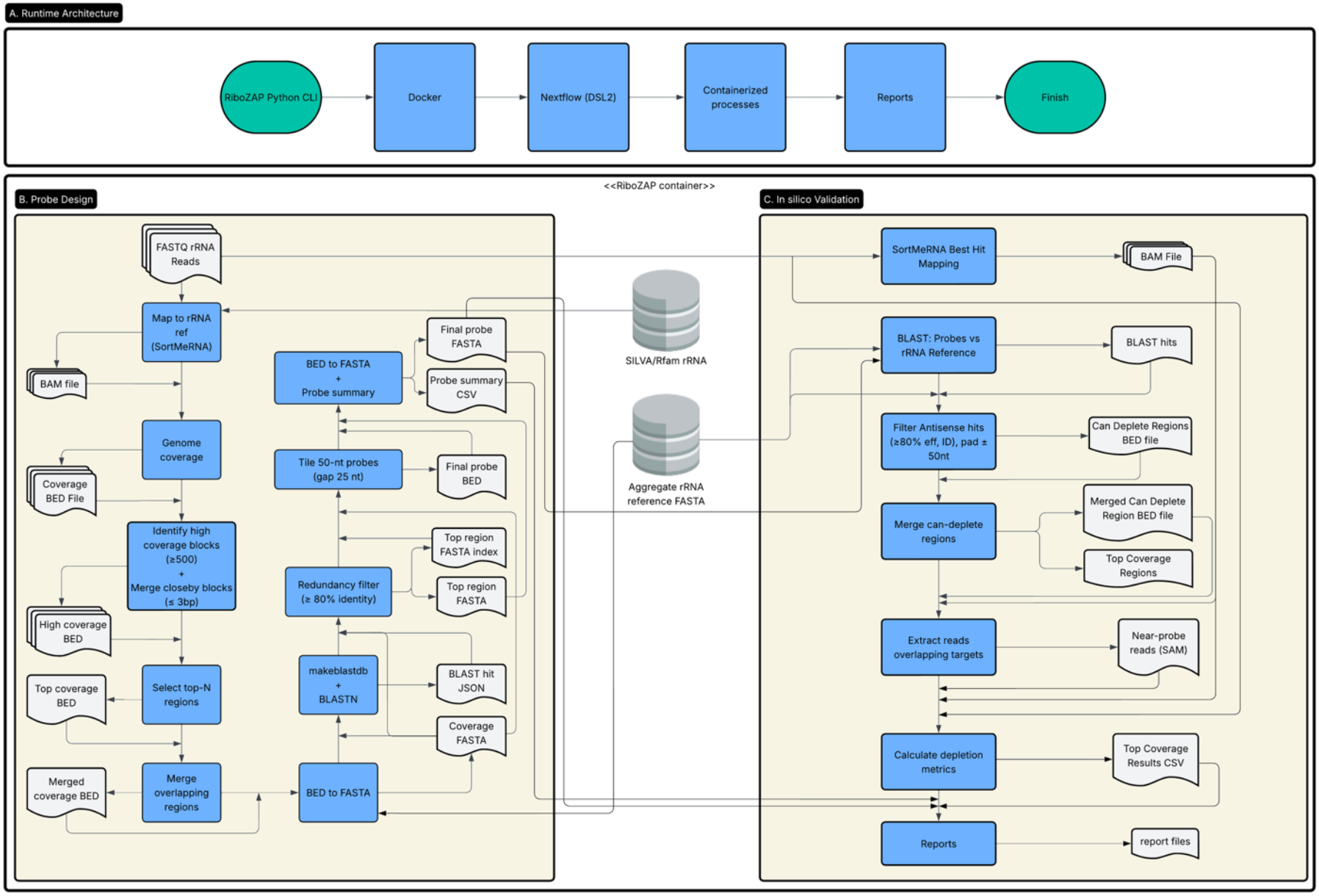
RiboZAP architecture and workflows. (A) Runtime orchestration via Python CLI, Docker and Nextflow. Dependencies and curated rRNA databases are bundled in the container.0020(B) Probe-design workflow: rRNA reads are aligned, coverage is summarized, high-coverage blocks are identified and merged, redundancy is removed by self-BLAST (≥80% identity), and 50-nt probes are tiled with a configurable 25-nt gap; outputs are final probes FASTA and a probe summary CSV. (C) In silico validation: Probes sequences are analyzed using BLASTN to identify can-deplete regions (≥80% effective identity, padded ±50-nt), best-hit rRNA reads are mapped, overlaps are counted, and depletion metrics are summarized in a CSV file along with a html report

### Probe design

The workflow begins with aligning rRNA-derived reads to curated rRNA references (SILVA and Rfam) using SortMeRNA. These alignments are processed with SAMtools to generate sorted, indexed BAM files, which provide the foundation for coverage analysis downstream. We then summarize per-base coverage with BEDTools (genomeCoverageBed -bga), producing compact BED intervals that capture stretches of uniform read depth across the reference.

From these coverage tracks, regions with strong rRNA signals are identified. High coverage blocks are defined as those exceeding a default threshold of 500 reads, and nearby blocks separated by only a few bases (≤3 bp) are merged to prevent biologically continuous targets from being split. Each block is then annotated with its length, maximum depth, and percentage of total reads mapped to the respective block. This allows prioritization of regions most likely to benefit from probe depletion. For each sample, the top N (default=50) blocks are retained and consolidated across datasets by merging overlaps, producing a non-redundant set of top rRNA coordinates.

From the top N retained blocks the sequences belonging to these intervals can be collected using BEDTools. Since rRNA regions can be highly conserved, the extracted sequences are compared against one another with BLASTN, and near-duplicates (≥80% identity) are collapsed to reduce redundancy. This allows the designed probes to have maximum specificity.

To obtain sufficient hybridization specificity and melting temp parameters, the length of the probes should be greater than 25-nt. To balance specificity, sensitivity and synthesis costs, 50-nt probes are recommended. However, longer probes will also perform adequately. The pipeline can be manually configured for a probe size of 25nt -100nt. The final output consists of a FASTA file containing all designed probe sequences, alongside a summary CSV table that consists of probe identifiers, sequences, and lengths.

An additional consideration is the total number of probes that can be included in the RiboZero Plus assay. The second enzymatic step of the protocol uses DNase to digest the DNA oligonucleotide probes; a critical step to limit the risk of left over depletion probes becoming included in the final RNAseq libraries. Too many probes will increase the probablity of saturating the ability of DNase to digest them. Based upon previous experience (data not shown) we recommend limiting the number of probes to <1500 at a concentration of 0.5 pmol per probe.

### In-silico Validation

Once the probe sequences have been designed, the next step in the pipeline is to validate whether they truly cover the rRNA regions targeted for removal. To do this, the new probes are first aligned back to the rRNA reference with BLASTN, which allows the assessment of how many probes map effectively to their intended targets. Resulting antisense alignments are then filtered using an effective identity threshold of ≥80% of the 50-nt probe length, and each retained hit is expanded by ±50 nt (default) to capture the realistic binding space. These filtered and padded BLAST hits are then merged to produce a non-redundant set of “can-deplete” rRNA regions that define the rRNA sequence space targeted by the probes. At the same time, a summary table is built to track coverage and provide a basis for downstream QC and reporting. Simultaneously, the original samples are re-aligned to the rRNA reference with SortMeRNA in best-hit mode (--num_alignments 1). This step is important because rRNA sequences are highly repetitive, sothis restrictioncan avoid artificially inflating the number of rRNA reads due to multi-mapping.

Once the depletable rRNA regions have been defined, the next step is to ask how many of the original sample reads fall into these regions. To do this, we extract from the BAM file all alignments that overlap the probe-targeted intervals using samtools view -L. This isolates the subset of rRNA reads that, in principle, would be removed by the designed probes. The reasoning here is: if a read maps within these intervals, then it lies in sequence space already covered by at least one probe and should be depletable in practice. By separating these reads into their own file, we can directly measure the size of this population and compare it against the total rRNA. This provides a clear, read-level view of probe efficiency and becomes the foundation for the depletion metrics downstream.

With probe-targeted reads isolated, the next step is to quantify probe performance at the sample level. For each dataset we record the total number of reads (T) from the FASTQ files, total mapped (M) as the number of alignment records to rRNA references, depleted (D) as the as the number of alignment records overlapping probe targeted regions, and unmapped (U = T-M), residual rRNA is (R = M - D).

Finally, all key outputs are compiled into a structured reports directory. This includes the final probe FASTA and summary tables, the depletion statistics, and a MultiQC report that aggregates metrics into a single view under “<outputdir>/probes” and “<outputdir>/reports” respectively. In the html reports, users can browse a probe design summary table that lists probe identifiers, sequences, and lengths. Sample-level mapping statistics are displayed as bar plots which illustrate the estimated depletion of rRNA, as well as the read composition per sample, broken down into depleted reads, remaining mapped rRNA, and unmapped reads. Together, these visualizations provide a glance at the overview of probe performance and depletion efficiency.

## Results

### In silico probe design and cross-sample validation

To evaluate the performance and cross-sample generalizability of RiboZAP-designed probes, we analyzed four mouse cecal MetaT datasets divided into two groups. The first group (Set 1; CC53 and CC54) was used to design the probes and to verify their performance on the same samples used for design, while the second group (Set 2; CC41 and CC63) served as an independent test set to assess performance on previously unseen data.

All datasets originated from RNA samples that had been experimentally depleted using the Illumina Ribo-Zero Plus Microbiome kit, which includes the Depletion Pool 1 (DP1) and Depletion Pool Microbiome (DPM) probe mixes as baseline rRNA removal reagents. Post-sequencing, FASTQ files were downsampled to 20 million total reads (10 million clusters) per sample using the Illumina BaseSpace Sequence Hub (BSSH) DRAGEN FASTQ Toolkit. To model further depletion achievable beyond this baseline, we computationally evaluated the supplemental probe sets generated by RiboZAP that correspond to the 2025 (20_25) and 5050 (50_50) experimental configurations previously tested in the laboratory (Roos et al., mSystems 2025).

These two configurations were selected from a broader probe-design matrix that varied both the number of high-coverage rRNA regions (20 to 50) and probe spacing (25 to 50 nt) to identify designs that maximized predicted depletion efficiency while minimizing probe count and synthesis cost. The 2025 configuration targeted the top 20 rRNA regions with a 25-nt tiling gap, producing 204 probes, whereas the 5050 configuration targeted the top 50 regions with a 50-nt gap, producing 206 probes.

As described in the implementation section, RiboZAP includes two workflows: one for probe design, which automatically performs in silico validation on the same samples used for design (CC53 and CC54), and another for standalone validation, which enables the newly generated probe sets to be evaluated independently on new datasets. For the cross-sample analysis in this study, we used the standalone validation workflow by providing the probe FASTA and summary files (--probes-fasta, --probes-summary) to test the designed probes against the non-design samples (CC41 and CC63). This approach allowed the same probe sequences designed from the training samples to be reproducibly tested on new data, allowing us to directly evaluate their depletion efficiency and consistency across biologically distinct mouse cecal samples.

### Probe performance on design and non-design samples

To understand how well the RiboZAP-designed probes perform in practice, we first tested them on the samples used for design (CC53* and CC54*). These samples provided a baseline to see how effectively the probes could recognize and remove the rRNA sequences from which they were designed. When applied to these datasets, the supplemental 2025 probe configuration achieved an average in silico rRNA reduction of 48% (Fig. 2A; Table S2). Read-composition analysis showed that most mapped rRNA reads overlapped probe-targeted regions, confirming strong coverage of the most highly expressed rRNA regions (Fig. 2B; Table S3). The two design samples showed nearly identical depletion patterns, indicating that RiboZAP’s probe-selection process performs consistently within the design set.

**Figure 2.**
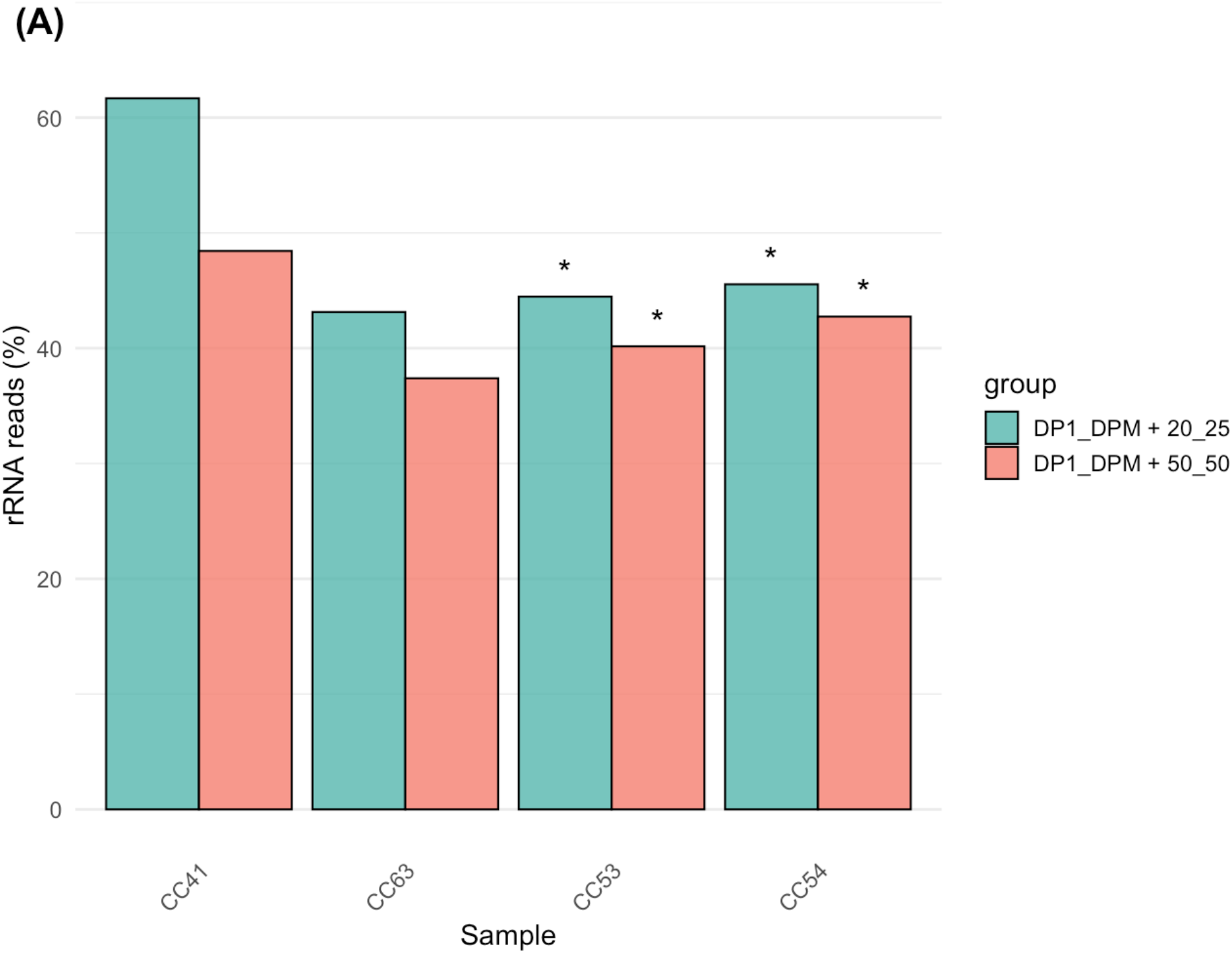
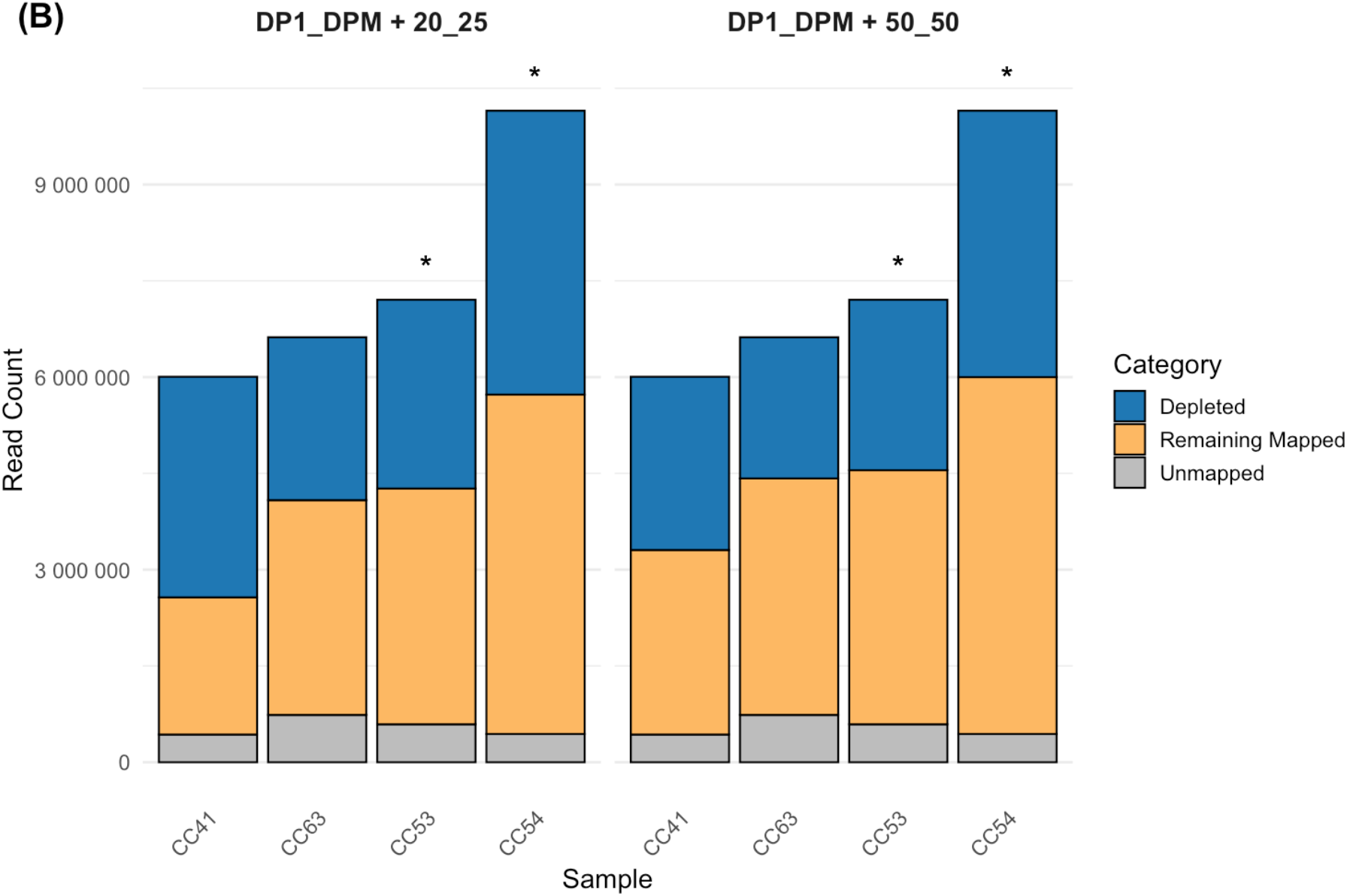
In silico evaluation and community-level impact of RiboZAP probe designs. **(A)** Predicted rRNA reduction percentages for design (CC53*, CC54*) and non-design (CC41, CC63) mouse cecal MetaT samples using the supplemental DP1_DPM+20_25 (top 20 rRNA regions, 25-nt gap) and DP1_DPM+50_50 (top 50 regions, 50-nt gap) probe configurations. Bars show consistent depletion across samples, with the DP1_DPM + 20_25 configuration predicted to remove an average of 48% of rRNA reads, compared to 42% with the DP1_DPM+50_50 configuration. **(B)** Read composition following simulated depletion. Stacked bars represent the proportions of unmapped (gray), remaining mapped (orange), and depleted (blue) reads for each sample, illustrating reproducible depletion patterns across both probe sets. Asterisks (*) denote samples used for probe design.

Next, we asked whether the same probes could work just as well in completely different samples that were not part of the design set (CC41 and CC63). Remarkably, the depletion efficiency in these non-design samples remained within a similar range (43-62%), showing that probes designed from one group of mouse cecal samples could still recognize and remove rRNA effectively in others. This consistency suggests that the rRNA profiles across individual mice share substantial sequence similarity, allowing a single probe design to generalize across multiple donors.

To explore how probe design parameters might influence depletion outcomes, we next evaluated the broader 5050 configuration, which targeted the top 50 rRNA regions with a 50-nucleotide tiling gap. This probe set achieved a moderate rRNA reduction of ∼42% across all samples (Fig. 2A; Table S2) while maintaining similar read composition profiles to those observed with the 2025 configuration (Fig. 2B; Table S3). Although slightly less efficient, the 5050 design still produced consistent depletion across both design and non-design samples, suggesting that distributing probes across more rRNA regions may reduce per-target efficiency but broaden coverage. Taken together, these results show that RiboZAP enables flexible probe design strategies that can be tuned to balance depletion efficiency, target coverage, and synthesis cost depending on experimental goals.

### Microbial diversity and Taxonomic composition of residual rRNA reads

To evaluate whether in silico depletion introduced any taxonomic bias, we analyzed the residual rRNA reads using the Illumina DRAGEN Metagenomics Pipeline Application on BaseSpace Sequence Hub to assess both microbial diversity and taxonomic composition. Shannon diversity indices were calculated from taxonomic profiles derived from residual rRNA reads (Fig. 3A; Table S1). Diversity values ranged from ∼3.0 to 4.5 across samples, with no significant differences between the 2025 and 5050 probe sets, indicating that the depletion process did not disproportionately target rRNA from specific taxa. A Wilcoxon rank-sum test confirmed that diversity differences between configurations were not statistically significant (*P = 0*.*69*), supporting the conclusion that probe-based depletion preserved the overall taxonomic balance. Together, these results demonstrate that RiboZAP can achieve effective rRNA reduction while maintaining community representation and minimizing the risk of probe-induced compositional bias.

**Figure 3.**
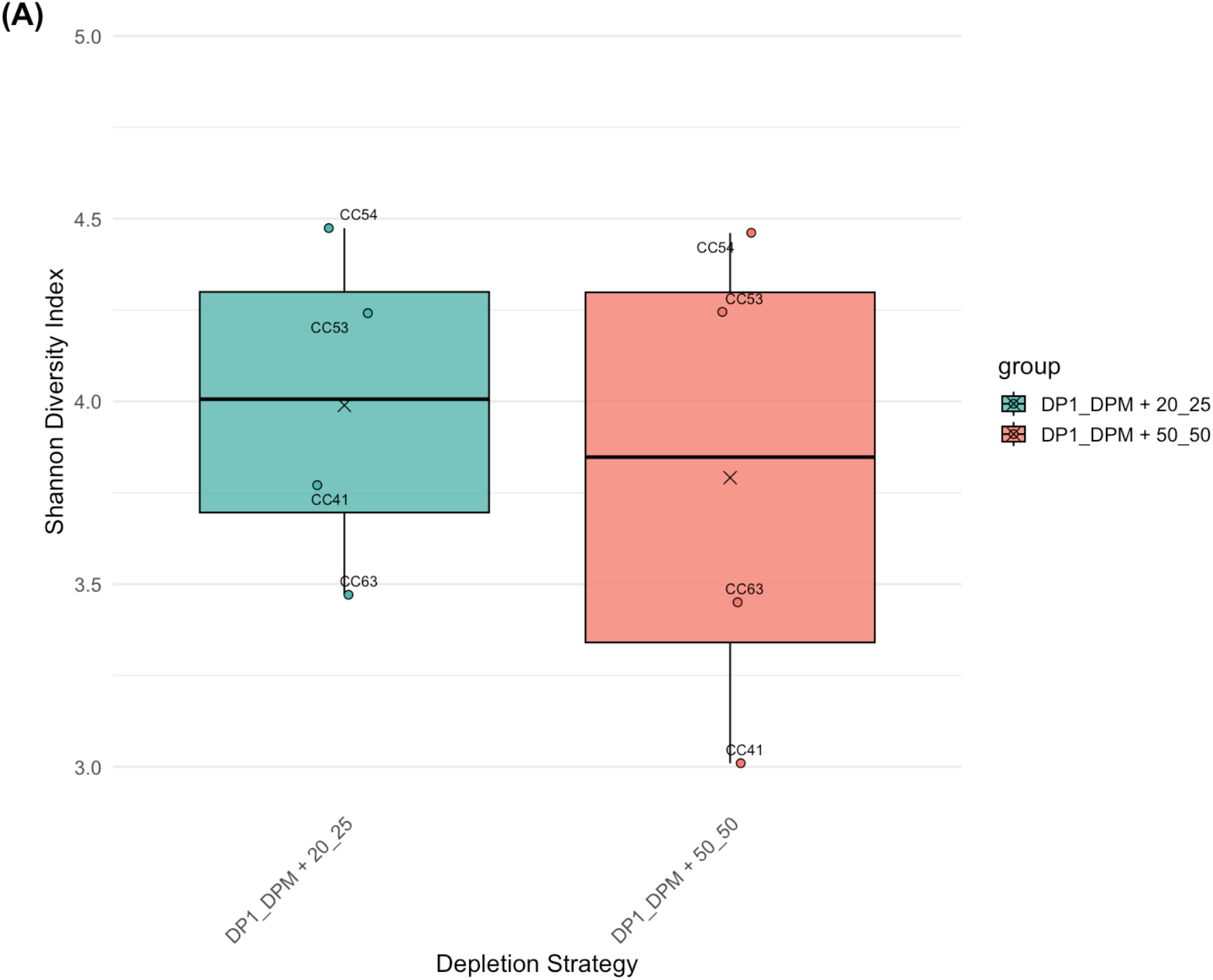
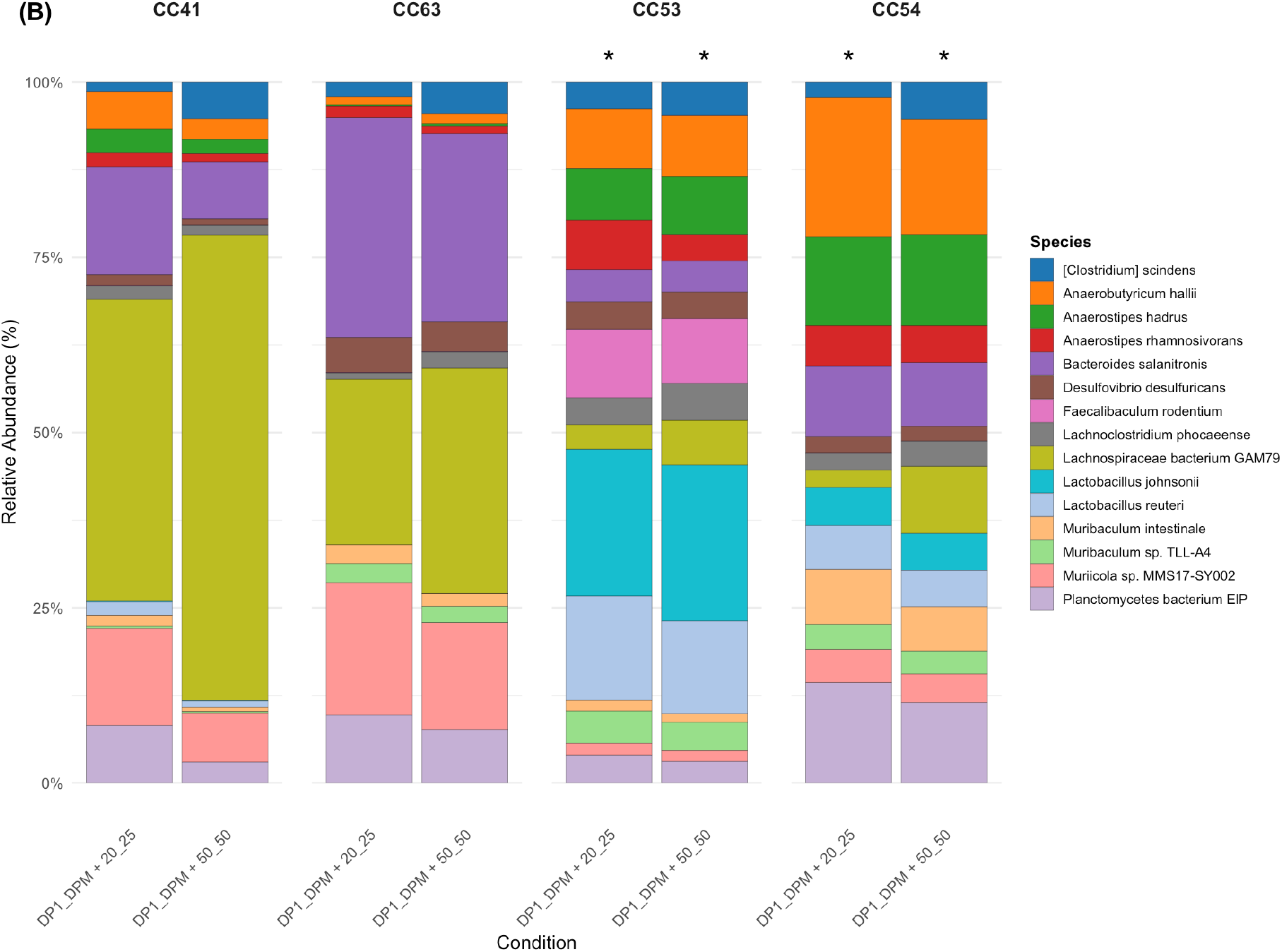
Microbial diversity and taxonomic composition of residual rRNA Metagenomics analyses were conducted using the Illumina DRAGEN Metagenomics Pipeline Application on BaseSpace Sequence Hub.(A) Microbial diversity analysis of residual rRNA reads. Shannon diversity indices were not significantly different between probe configurations (P = 0.69; Wilcoxon rank-sum test), confirming that rRNA depletion did not bias microbial community structure. (B) Taxonomic composition of residual rRNA reads following simulated depletion. Bars represent the total abundance across design (CC53*, CC54*) and non-design (CC41, CC63) samples. The even distribution of taxa across conditions indicates that RiboZAP-based rRNA depletion did not preferentially remove or enrich rRNA from specific species, thereby preserving the overall microbial community structure. Asterisks (*) denote samples used for probe design.

To further examine potential taxonomic bias following in silico depletion, the top microbial taxa contributing to the residual rRNA fraction were identified (Fig. 3B). Relative abundance profiles showed that no single species dominated after simulated depletion, and the overall community composition remained evenly distributed across both design and non-design samples. The top 15 most abundant species were consistently represented under both the 20_25 and 50_50 probe configurations, confirming that probe targeting was broad and not species-restricted (Fig. 3B; see Fig. S1 for the top 100 species). These results indicate that RiboZAP effectively depletes rRNA while preserving microbial community structure.

### Comparison with experimental validation

The *in silico* predictions obtained in this study closely reflected the experimental results previously reported [3]. In that work, supplemental RiboZAP-designed probe sets improved rRNA removal across all mouse cecal samples tested, increasing the proportion of retained mRNA-rich reads from roughly 60% to 75% (*P < 0*.*01*). This mirrors the depletion efficiencies predicted computationally, indicating that the *in silico* modeling accurately captured real experimental performance. Importantly, both approaches produced consistent taxonomic profiles, with no evidence of depletion bias toward particular microbial or host transcripts. The strong agreement between computational and laboratory observations highlights RiboZAP’s predictive reliability and demonstrates its value as a design tool for optimizing rRNA depletion strategies before costly wet-lab experiments.

## Conclusion

In this study, we demonstrated that RiboZAP can accurately model and predict rRNA depletion performance using an *in silico* framework. Probe sets designed from a small number of mouse cecal MetaT samples performed consistently when applied to additional, non-design samples, indicating that the approach is broadly transferable across related datasets. The close agreement between *in silico* predictions and experimental results reported previously [3] supports the accuracy of RiboZAP’s design algorithm. By allowing researchers to test and refine probe sets computationally before synthesis, RiboZAP provides a practical way to improve rRNA removal strategies, reduce experimental costs, and accelerate the development of custom probe designs for complex microbial communities.

## Data and code availability

All the code is available on GitHub at https://github.com/ilmn-mouse-cecal/ribozap. The FASTQ files obtained for this work are available via the SRA under BioProject accession ID PRJNA1258316.

