## supplementary file 1 for "RiboZAP: Custom probe design for rRNA depletion in complex metatranscriptomes" (from another version of this preprint: version 1, 2025-11-07)

|  | **sample_name** | **group** | **shannon_index** |
| --- | --- | --- | --- |
| **1** | CC41 | DP1_DPM + 20_25 | 3.77078 |
| **2** | CC41 | DP1_DPM + 50_50 | 3.00919 |
| **3** | CC53 | DP1_DPM + 20_25 | 4.24155 |
| **4** | CC53 | DP1_DPM + 50_50 | 4.24449 |
| **5** | CC54 | DP1_DPM + 20_25 | 4.47457 |
| **6** | CC54 | DP1_DPM + 50_50 | 4.46108 |
| **7** | CC63 | DP1_DPM + 20_25 | 3.47064 |
| **8** | CC63 | DP1_DPM + 50_50 | 3.45054 |

Table S1. Shannon diversity indices calculated from residual rRNA reads following in silico depletion across design (CC53, CC54) and non-design (CC41, CC63) mouse cecal metatranscriptomic samples. Shannon diversity values were derived from species-level taxonomic profiles generated using the Illumina DRAGEN Metagenomics Pipeline on BaseSpace Sequence Hub.

| **Sample** | **rRNA Depletion Percent** | **group** |
| --- | --- | --- |
| **CC54** | 45.54 | DP1_DPM + 20_25 |
| **CC53** | 44.47 | DP1_DPM + 20_25 |
| **CC54** | 42.74 | DP1_DPM + 50_50 |
| **CC53** | 40.16 | DP1_DPM + 50_50 |
| **CC63** | 43.14 | DP1_DPM + 20_25 |
| **CC41** | 61.67 | DP1_DPM + 20_25 |
| **CC63** | 37.39 | DP1_DPM + 50_50 |
| **CC41** | 48.43 | DP1_DPM + 50_50 |

Table S2. Predicted rRNA depletion percentages across design (CC53, CC54) and non-design (CC41, CC63) mouse cecal metatranscriptomic samples. Values represent the proportion of rRNA reads predicted to be removed following in silico depletion using RiboZAP supplemental probe sets DP1_DPM + 20_25 and DP1_DPM + 50_50. Percentages were calculated from the top_coverage_result.csv output generated by the RiboZAP validation workflow.

| **Sample** | **Unmapped** | **Remaining Mapped** | **Depleted** | **group** |
| --- | --- | --- | --- | --- |
| **CC54** | 440066 | 5287556 | 4422088 | DP1_DPM + 20_25 |
| **CC53** | 591170 | 3672619 | 2941233 | DP1_DPM + 20_25 |
| **CC54** | 440066 | 5559797 | 4149847 | DP1_DPM + 50_50 |
| **CC53** | 591170 | 3957923 | 2655929 | DP1_DPM + 50_50 |
| **CC63** | 737395 | 3345638 | 2538179 | DP1_DPM + 20_25 |
| **CC41** | 432024 | 2136562 | 3436946 | DP1_DPM + 20_25 |
| **CC63** | 737395 | 3683570 | 2200247 | DP1_DPM + 50_50 |
| **CC41** | 432024 | 2874207 | 2699301 | DP1_DPM + 50_50 |

Table S3. Read composition summary following in silico rRNA depletion across design (CC53, CC54) and non-design (CC41, CC63) samples. Counts represent the number of reads classified as unmapped, remaining mapped, and depleted after applying the RiboZAP supplemental probe configurations (DP1_DPM + 20_25 and DP1_DPM + 50_50). Values were derived from the top_coverage_result.csv output of the RiboZAP validation workflow and correspond to the data visualized in Figure 2B.

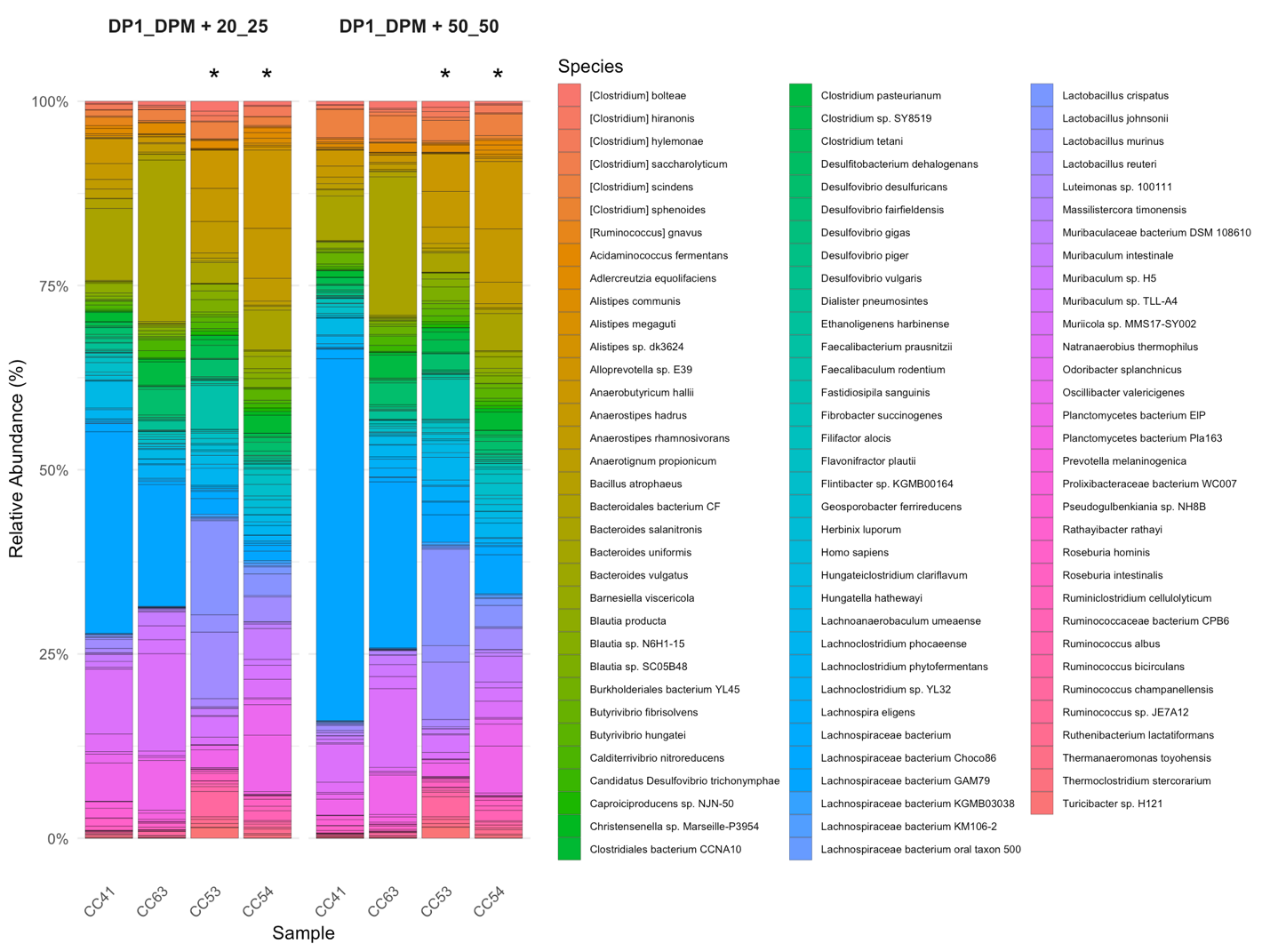

Figure S1. Relative abundance of the top 100 microbial species detected in residual rRNA reads following in silico depletion across design and non-design samples. Species-level abundances were derived from the microbial-classification-report.tsv output of the Illumina DRAGEN Metagenomics Pipeline on BaseSpace Sequence Hub. The top 100 species were identified by summing read counts across all samples and selecting the taxa with the highest total abundance. Both DP1_DPM + 20_25 and DP1_DPM + 50_50 probe configurations produced evenly distributed residual rRNA profiles, with no single species dominating after depletion.
